# The Role of the Medial Prefrontal Cortex in Spatial Margin of Safety Calculations

**DOI:** 10.1101/2020.06.05.137075

**Authors:** Song Qi, Logan Cross, Toby Wise, Xin Sui, John O’Doherty, Dean Mobbs

## Abstract

Humans, like many other animals, pre-empt danger by moving to locations that maximize their success at escaping future threats. We test the idea that spatial margin of safety (MOS) decisions, a form of pre-emptive avoidance, results in participants placing themselves closer to safer locations when facing more unpredictable threats. Using multivariate pattern analysis on fMRI data collected while subjects engaged in MOS decisions with varying attack location predictability, we show that while the hippocampus encodes MOS decisions across all types of threat, a vmPFC anterior-posterior gradient tracked threat predictability. The posterior vmPFC encoded for more unpredictable threat and showed functional coupling with the amygdala and hippocampus. Conversely, the anterior vmPFC was more active for the more predictable attacks and showed coupling with the striatum. Our findings suggest that when pre-empting danger, the anterior vmPFC may provide a safety signal, possibly via predictable positive outcomes, while the posterior vmPFC drives prospective danger signals.

---

Staying in close proximity to safety is a key antipredator behavior as it increases the likelihood of the organism’s future escape successs^1^. One metric used by behavioral ecologists to measure this safety behavior is called spatial margin of safety, where prey will adopt locations that prevent lethal predatory attack^1–3^. In turn, this provides the prey with a safety net, while also reducing stress, energy consumption and promotes increased focus on other survival behaviors, such as foraging and copulation. Humans appear to use safety distance in similar ways. For example, when human subjects are placed close to a safety exit, measures of fear decrease and when under threat, and the sight of safety signals reduces fear and fear reinstatement^5–7^. Here, we test the idea that when subjects are pre-empting threats of varying attack location probabilities, subjects will vary their spatial margin of safety (MOS) decisions depending on predictability. We propose that MOS decisions involve prospective spatial planning, which involves estimating safety by calculating the predator’s attack locations^4^. Further, we examine how pre-emptive MOS decisions are instantiated in human defensive circuits^9,10^.

In the natural world, prey encounter predators that attack with varying degrees of uncertainty. Uncertainty is often determined by the likelihood of attack and the distribution of distances at which the threat will attack. For example, uncertainty alerts the prey that information about the predator’s impending attack location is unknown, thereby resulting in increased anxiety and movement towards safety^5^. Thus, pre-empting predation via close spatial MOS, safeguards against the unpredictable spatial and temporal movements of the predator^6^. Consequently, the ability to predict a predator’s attack location will in turn shape the prey’s MOS calculations, whereby uncertain threats will result in low risk behaviors and smaller spatial radius from a refuge at the expense of forgoing other survival needs (e.g. food). In particular, frequent and salient outlier information in a given information, as presented as leptokurtic noise, makes organisms prone to overreaction and inaccurate estimations of the environment^13^. Therefore, our second question is how statistical uncertainty of a threat’s attack location sways spatial MOS decisions and shifts activity in the human defensive circuits.

The prospective nature of MOS decisions may elicit activity in a set of neural circuits involved in anxiety^7^, which can be defined as a future oriented emotional state and involves the behavioral avoidance of potential dangers. Two drivers in this spatial avoidance are the ventromedial prefrontal cortex (vmPFC), and the hippocampus^7–10^. For example, the hippocampus plays a key role in anxiety, and guides decisions via memory and prospection^18,19^. Further, synchronization between the hippocampus and vmPFC are associated anxiety like behaviors^20–22^, suggesting that the hippocampus, potentially along with the amygdala, is involved in signaling the threat significance of a stimulus. The vmPFC is a heterogeneous structure involved in information seeking, anticipation and the organization of defensive and safety responses^11–14^. Research has shown that a safety stimulus during an aversive experience results in increased activity in the anterior vmPFC while decreasing threat also results in increased activity in the same region, suggesting that the anterior vmPFC may emit safety signals^5,26^. Research also shows that attention set to safety signals, extinction, and down-regulation of anxiety are associated with vmPFC activity, suggesting that it is a key node in what has been called the fear suppression circuit^27–29^. Conversely, the posterior vmPFC, encompassing the subgenual and rostral anterior cingulate cortex (sgACC and rACC), receives dense projections from the amygdala^15^ and is implicated in negative affective responses and behavioral expression of fear,^11,16,31,32^. How these, and other brain regions are evoked during pre-emptive MOS decisions is yet to be tested.

To address these gaps in knowledge between spatial MOS decisions and human defensive circuits, we created a task to investigate spatial MOS decisions under uncertainty and elucidate: (i) How do changes in the threat’s attack predictability, threat intensity, and reward value impact the subjects’ MOS decisions? And ii) Do the hippocampus and vmPFC encode characteristics of threats that are central to MOS decisions? This task models the ecological phenomena where animals venture further away from their safety refuge to acquire adequate supplies of food. To create less predictable attack positions, we used leptokurtic distributions, which are evolutionarily novel and volatile in nature, and have been shown to increase the level of uncertainty and difficulty to learn to the environment^13^. Leptokurtic noise is generated as the composite of two normal distributions with similar means and contrasting variances. Leptokurtic distributions are thus probability density curves that have higher peaks at the mean and are fatter tailed where extreme outcomes (outliers) are expected more (Fig. C). We contrasted this with standard Gaussians (Fig. 1D and E)), which are more computationally familiar. We hypothesized that when subjects are facing virtual predators with higher frequency of outlier attack distributions, this will result in more uncertainty and therefore, decisions to move closer to safety.

**Fig. 1.**
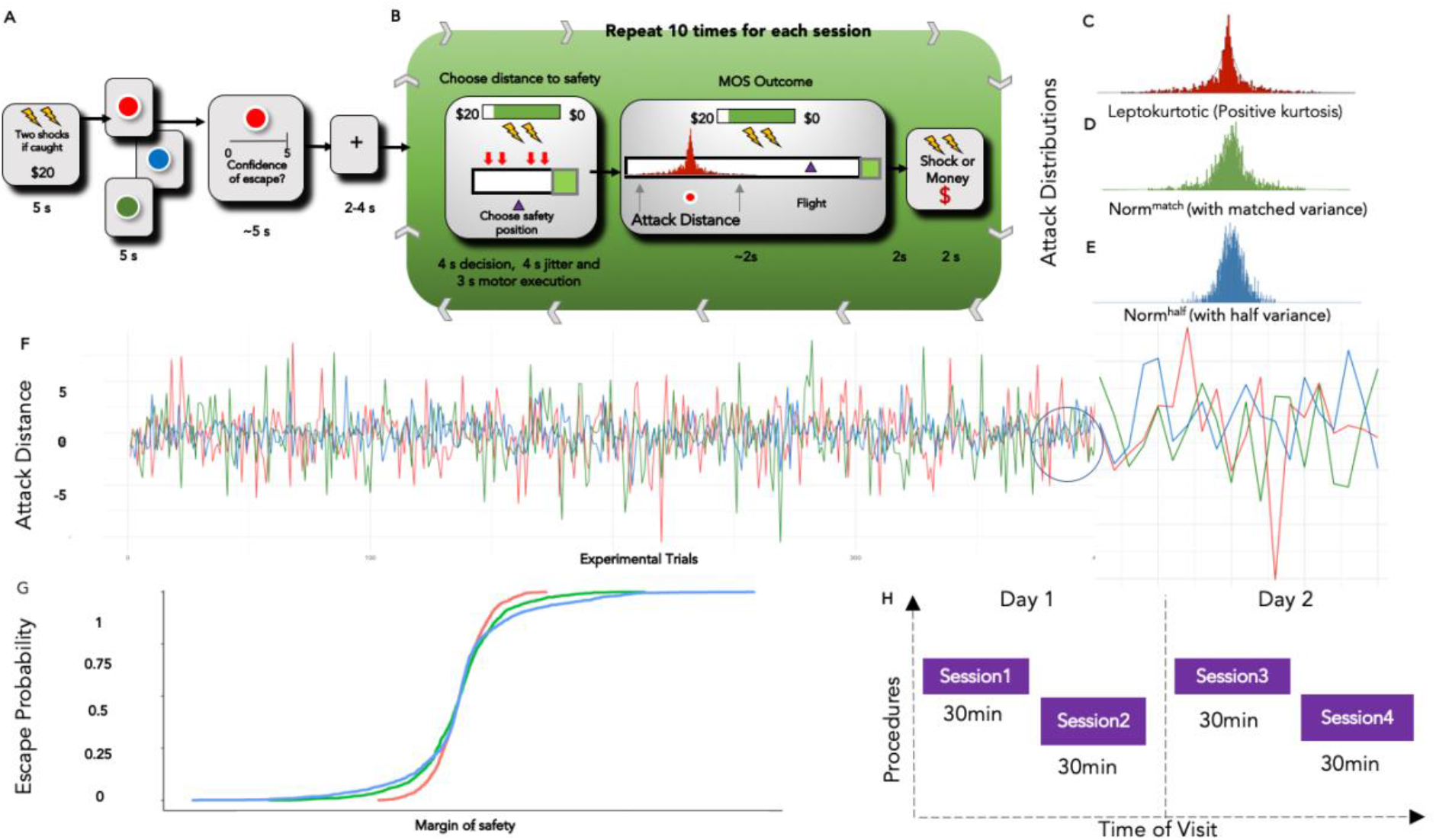
Experimental structure. (A) During the MOS decision task, participants were first presented with a series of information screens at the beginning of every 10 trials (one trial block), displaying the reward/shock level, color of the predator (leptokurtic,where a kurtosis is added to the normal distribution resulting in heavy tails -red, norm_match_, where the variance is matched with the leptokurtic condition-green, norm_half_, where the variance is half as compared to the leptokurtic condition - blue), before being asked to rate their confidence in escaping. In the low shock/reward conditions, participants receive 1 shock and the base reward respectively. In the high shock/reward conditions, participants received 2 shocks and twice the base reward. (B) On every trial, for the first 4 seconds, participants were presented with a screen displaying the margin of safety runway and their initial location. They were told to make a choice of where to put themselves during this phase. However, they were not able to actually move in this phase. After a 4 second jitter, they were presented with the same screen again where they can move to the desired MOS with a varying bar of reward meter depending on the MOS location. In the next 2 seconds, the outcome of the chasing was revealed, including whether their escape was successful and how much reward was gained. (C) Attack distributions for leptokurtic volatile; (D) gaussian distribution with matched variance and (E) half the variance gaussian; (F) the predator’s attack distances through all trials. Zero on the Y axis marks the mean of the distribution, while numbers represent how far away the drawn instance is away from the mean. (G) Escape probability. X axis represents possible margin of safety choices, while Y access represents the corresponding probability of escape. (H) Schematic representation of the experimental procedure. Participants undergo 4 x 30 min scans sessions over a two-day period.

## Results

### MOS choices are less risky in the less predictable threat environment

MOS choice in the task represents the position participants selected relative to the safety refuge. In order to investigate how uncertainty of predator attack modulates MOS choices, we first examined how MOS decisions varies across distributions types, with a repeated-measures, one-way ANOVA to investigate how MOS decisions varied across distribution types. The result showed a main effect of distribution type [F(2,44) = 61.33, p < 0.001]. A Tukey post hoc test revealed that participants’ MOS choice was significantly closer to the safety zone in the leptokurtic distribution condition (0.74 + −0.06) than in the norm_match_ condition (0.68 +−0.03) and norm_half_ condition (0.67 +−0.01). This indicates that participants perceived leptokurtic distributions as more risky, resulting in overall safer choices. Interestingly, there was no significant difference in mean MOS choices between the two normal distributions. This suggests that only a fundamental change in the statistical structure of the target distribution can impact participants’ decision under threat, rather than a change in the variance of the distribution (Figure 2 a,b,c,d)

**Fig. 2.**
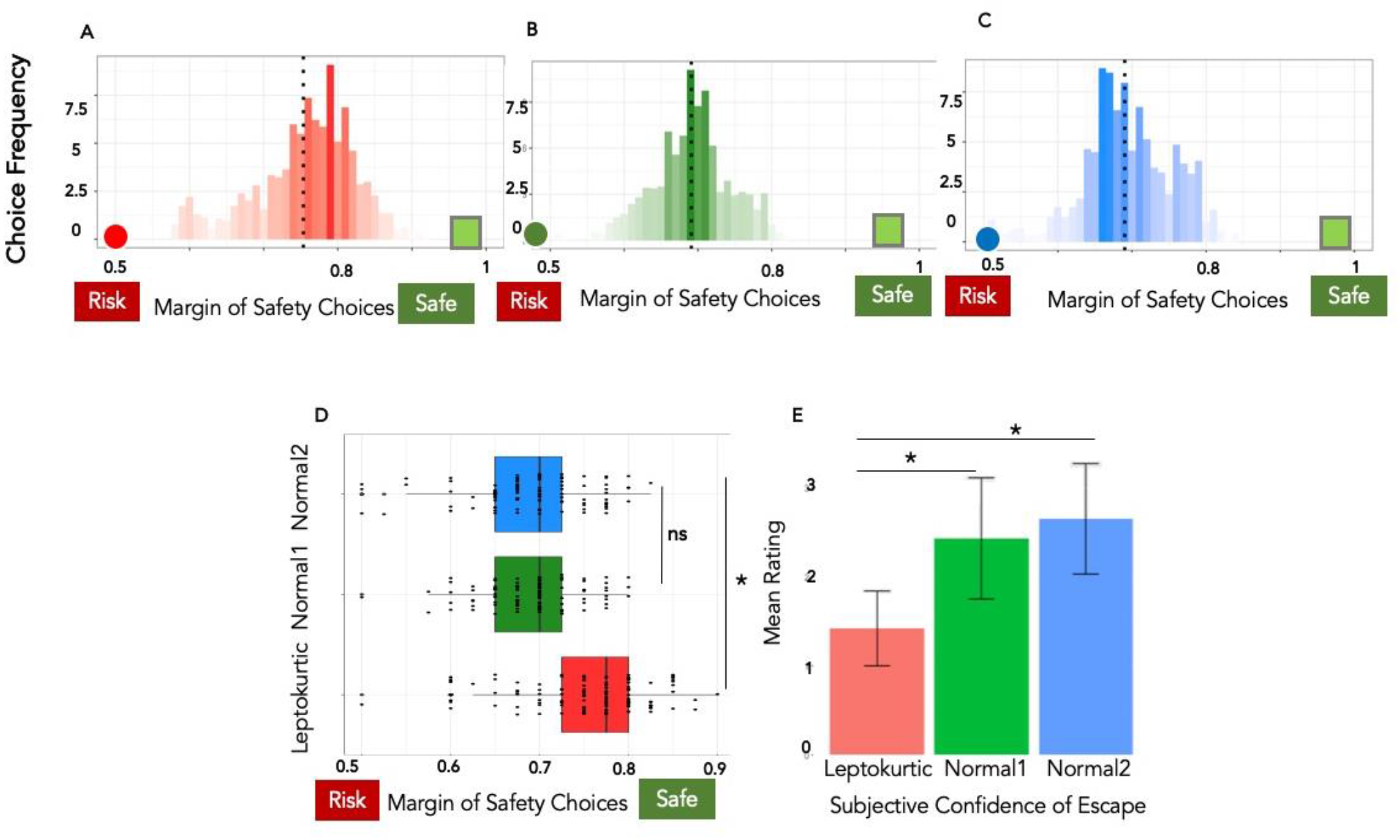
Behavioral Results. Choice frequencies for (A) leptokurtic, (B) matched variance and (C) half variance attacking threats. The MOS decision phase and the outcome. (D): Confidence ratings for leptokurtic distribution, matched variance normal distribution, and normal distribution with half variance. Post-hoc analysis revealed that participants were less confident in the leptokurtic condition compared to the other two conditions (p < 0.001). Leptokurtic attack location are in red; normal distribution with matching variance are in green; and normal distribution with half variance are in blue.

### MOS choices are less risky in threat environment with higher punishment

To further disentangle how shock and reward levels could interact with predator attack type as additional external incentives, we examined participants’ MOS choices within high/low shock conditions and high/low reward conditions. In the low shock/reward conditions, participants receive 1 shock and the base reward respectively, whereas in the high shock/reward conditions, participants received 2 shocks and twice the base reward. While there was no significant difference in their MOS decisions when facing different levels of rewards (t(21) = 1.378, p = 0.182) their MOS choices were significantly more conservative in the high shock condition (0.75 +−0.07), compared to the low shock condition (0.69+-0.05): t(21) = 21.21, p < 0.001. This suggests that participants were sensitive to the level of danger and adjusted their MOS decisions accordingly (Supplementary figure 1).

### More confident participants made riskier MOS decisions

Having shown that uncertainty in the attack distribution influences observable behavior, we asked whether it also affects subjective confidence in escape success. We collected participants’ confidence ratings before every unique trial block (shown in figure 1 A/B, every 10 trials consist a unique trial block). An ANOVA on the confidence ratings also revealed that participants were generally more confident on trials in the two normal distributions compared with trials in the leptokurtic distribution. A main effect of distribution type was found [F(2,44) = 27.32, p < 0.001], and a Tukey post hoc test showed that confidence rating in the leptokurtic condition (1.42 +−0.42) was significantly lower than those in the norm_match_ condition (2.43 +−0.68) and the norm_half_ variance (2.65 +−0.62) (p < 0.001) (figure 2 e). We also examined the relationship between participants’ MOS choices and confidence ratings. Interestingly, a significant correlation was only observed in the leptokurtic condition, where individuals who were more confident made riskier MOS choices (r = −0.54. p = 0.04). This effect was not observed for either the norm_match_ condition (r = 0.25, p = 0.37) nor the norm_half_ condition (r = −0.31, p = 0.27).

### MOS decisions are represented within prefrontal and subcortical regions

Building on our behavioral results, we next sought to identify neural systems underlying MOS decisions in response to varying levels of attack uncertainty. Due to the design feature of the behavioral experiment, the decision phase consists of both a cognitive (perception of the threat) and decision component, making the univariate analysis insufficient to capture the underlying dynamics of the neural process^34,35^. The MVPA analysis here thus serves two main purposes: to identify the key regions involved in decision making under the current threat, and to distinguish the underlying neural mechanism among threats with different levels of uncertainty. Results of this analysis can then be used to inform ROIs for subsequent connectivity and parametric modulation analysis. To accomplish this, we used a searchlight cross-decoding approach using linear support vector regression (SVR) and leave-one-out cross-validation (Supplementary Methods).

**Fig. 3.**
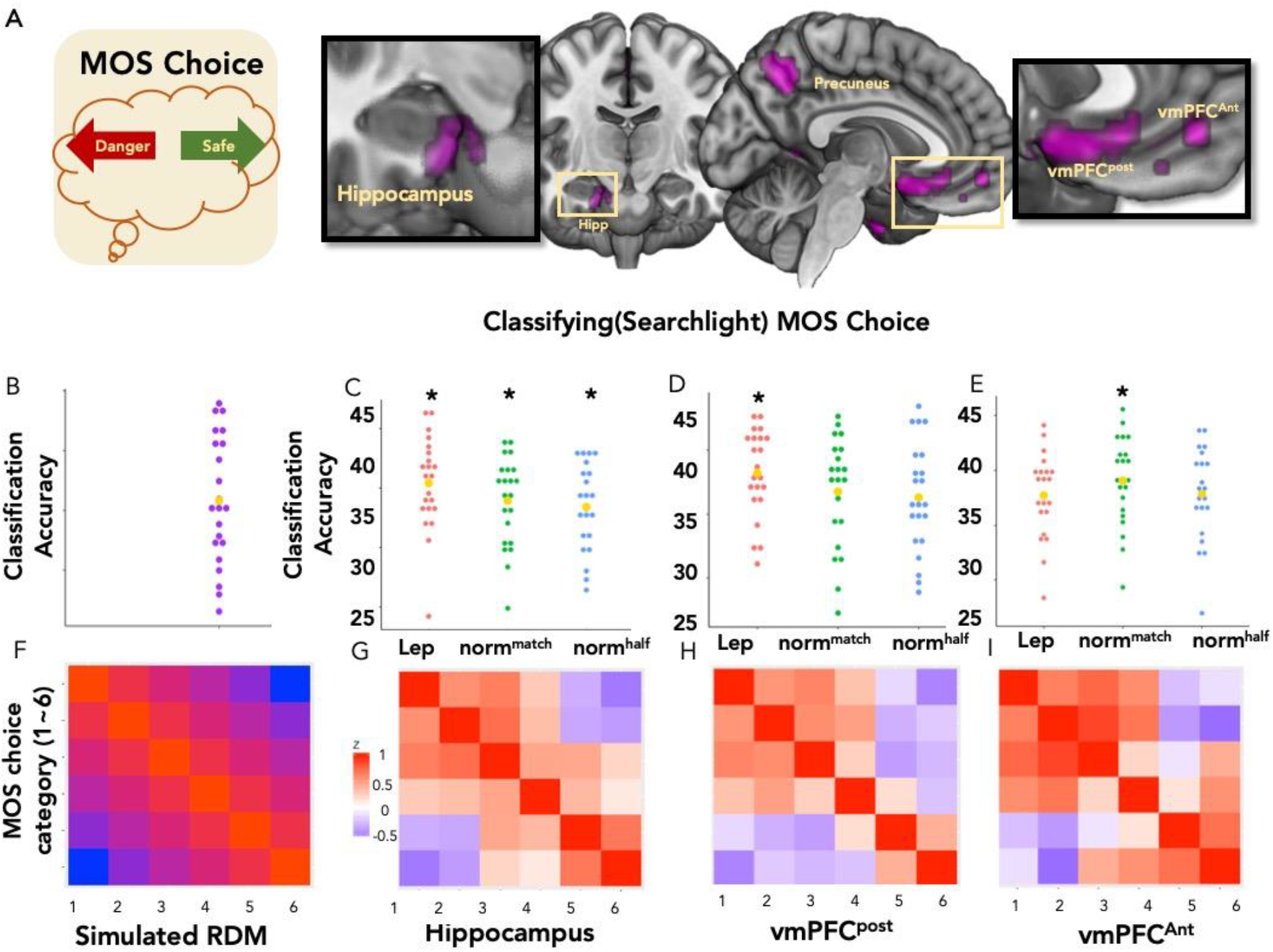
Neural representation of pre-emptive MOS decisions. Avoidance decisions decoded in the vmPFC, Hippocampus. (A): whole brain search light map displaying regions significantly activated for the MOS choice classifier (FDR corrected, p < 0.05). (B): Classification accuracy of the MOS choice classifier. Each dot represents data from a single subject. Average accuracy was significantly higher than the simulated chance level (p < 0.001). Box and whisker plots display accuracies from the same classifier, at various regions of interest (the hippocampus, vmPFC (posterior) and vmPFC (anterior)).(C): In hippocampus, classification accuracy from all three attack conditions were significantly higher than their corresponding chance levels. (D); Classification accuracy was only significantly higher than chance in the leptokurtic distribution in vmPFC_post_.(E): Classification accuracy was only significantly higher than chance in the norm_match_ condition in vmPFC_ant_. (F): Behavioral similarity structure among MOS choices. The Behavioral similarity structure represent how similar MOS choices are on the behavior level. For example, MOS choice 1 and 2 are closer in distance compare to choice 1 and 6, thus more similar in the structure. Naturally, choices are more similar when in close spatial distance, and more dissimilar when in sparse spatial distance. (G): Actual pattern similarity within the regions of interest. The neural RDM in the hippocampus was significantly correlated with the theoretical model (*r* = 0.593, *P* < 0.001). Similar correlation effects were also found in (H) vmPFC_post_ and (I) vmPFC_ant_, (r = 0.754, p < 0.001; r = 0.482. p < 0.001).

Two separate whole brain searchlight analyses were performed to answer the following questions respectively: which regions are critically involved in processing 1) different attacking distributions and 2) Margin of safety choices. The first classifier predicted which distribution type a given trial belonged to. This showed that regions including right insula and the mid-cingulate cortex (MCC) encoded the distribution type, with a decoding accuracy significantly higher than the Monte-Carlo simulated chance level accuracy (overall accuracy: t(21) = 2.82, p =.010). The whole brain decoding map was thresholded at P<0.05 (FWE) (Fig. 3a).

Next, for the analysis of MOS decision types, each trial was labelled according to the MOS decision the subject made, and a classifier was trained to predict which trials fall into which decision categories. The categories were created by grouping MOS choices that are close in spatial distance together. i.e. the entire MOS choice runway is divided to six segments from left to right). Each choice category thus represents a level of how close participants place themselves to the safety. Decoding of choices was found in regions including the right hippocampus, vmPFC_post_ and vmPFC_ant_ with a decoding accuracy significantly higher than chance level (t(21) = 2.47, p = .022). These results suggested that the both the distribution type and MOS decision making process is robustly represented in the above mentioned regions. (Fig. 3 A,B)

### vmPFC subregions differentially encode MOS decisions according to uncertainty

The regions implicated in the whole brain searchlight overlap with ROIs in previous literature shown to be critically involved in the process of decision making under threat. We thus performed MVPA analysis within each ROI, namely the hippocampus, vmPFC_post_ and vmPFC_ant_ to investigate how they uniquely contributed to the MOS decision process. Within each specified ROI, we investigated classification accuracy for the MOS decisions labels, separately for each distribution conditions. Thus, by comparing how well the process is decoded within each ROI, we can examine how the involved regions drive behavioral change depending on levels of uncertainty in different predator conditions.

Within the vmPFC_pos_, only choice decoding for the leptokurtic condition was significantly above the Monte-Carlo simulated chance level (Monte-Carlo simulated baselines: leptokurtic, 36.7%; norm_match_, 34.8%; norm_half_, 33.7%) (leptokurtic distribution, p < .001; norm_match_, p =.410; norm_half_, p = .868). Within the vmPFC_ant_, only classification for the norm_match_ condition was significantly above chance level (leptokurtic distribution, p = .341; norm_match_, p =.004; norm_half_, p = .156). Within the hippocampus, classification for all 3 distribution types was significantly above chance level (leptokurtic distribution, p < .001; norm_match_, p = .011; norm_half_, p = .038). A follow up ANOVA did not reveal a significant difference among the decoding accuracies (Fig. 3 B,C,D,E).

### Univariate overlap with vmPFC regions involved in ‘fear’ and extinction

To compare the activated regions with past studies, we constructed ROIs from neurosynth using the key words “fear” (for comparison with posterior vmPFC/sgACC) and “extinction” (for comparison with vmPFC_ant_ ROIs were constructed using 6mm spheres from the peak coordinate. Small volume corrections were performed. Extinction maps were used as we hypothesized that the extinction and reduced threat would overlap. We then performed SVC with the “fear” ROI on vmPFC_pos_ with the leptokurtic contrast (p < 0.001, T = 5.07, cluster size = 31, (0,26,−12)) and SVC with the “extinction” ROI on vmPFC_ant_ (p = 0.010, T = 4.35, cluster size = =11, (−2,46,−10)). For a full list of activated regions, please refer to supplementary table 1. These coordinates overlap with the corresponding ROIs taken from the searchlight analysis, indicating that information processing and learning through both fear and safety are potentially presented in MOS decision making through vmPFC_post_ and vmPFC_ant_, respectively.

### vmPFC activity encodes MOS decisions

Having demonstrated that vmPFC activity patterns encode MOS decisions, the next step was to ask whether overall BOLD activity levels in the vmPFC also covaried with MOS decision (Fig. 4E). To test this, we constructed two univariate parametric modulators indicating whether the participants’ final MOS choices is a safety choice or a risky choice (compared to their randomly assigned initial location). The parametric modulation of univariate data thus reveals what regions showed activity associated with risky/safety choices under different levels of predictability. Inspection of the resulting statistical maps, using SVCs from the previously constructed vmPFC_post_ and vmPFC_ant_ ROIs, showed that the “move to danger” and “move to safety” modulations were significant in the vmPFC_post_ and vmPFC_ant_ ROIs respectively (Move to danger: p < 0.001, T = 6.44; Move to safety: p < 0.001, T = 4.39, supplementary table 4).

### Representational similarity analysis of the vmPFC_post_, vmPFC_ant_ and hippocampus

The MVPA searchlight analysis offers insights into what key regions are involved in coding MOS decision process. However, it is left unclear how different MOS choices were actually neurally represented in the ROIs mentioned above. Thus, a representational similarity analysis was conducted to investigate the underlying geometry of the neural encoding of the MOS decision variables in the above mentioned ROIs. Distinctive clustering in the RDM structure also help further validate the original behavioral paradigm, showing how sensitive participants were to all the possible MOS choices.

A Behavioral RDM, together with RDMs from the neural data within the hippocampus, vmPFC_post_, and vmPFC_ant_ were constructed to investigate the potential MOS decision information and perceived distribution information embedded in the activity patterns of these ROIs. A high level of similarity between the theoretical structure and the actual brain activity in a certain ROI will indicate that task-relevant information is encoded in a way that is consistent with the behavioral structure of the during the MOS decision process. Figure 3 illustrates the theoretical/behavioral RDMs constructed by the pairwise relations of the 50 MOS decisions. Spearman correlation coefficients were used to calculate the distance between the model and neural data matrices. The neural RDM in the hippocampus was significantly correlated with the theoretical model (*r* = 0.593, *P* < 0.001) across all conditions. Similar correlation effects were also found in vmPFC_post_ and vmPFC_ant_, (r = 0.754, p < 0.001; r = 0.482. p < 0.001), but these were specific to the leptokurtic and norm_match_ conditions respectively (fig 3f,g,h,i).

Converging evidence from the previously mentioned searchlight, univariate parametric modulation, and RSA analysis has shown that the vmPFC subregions (vmPFC_post_ and vmPFC_ant_) play a vital role in the encoding of MOS decisions under environments with different levels of predictability. Next we further investigate the connectivity structure seeding from these regions.

**Fig. 4.**
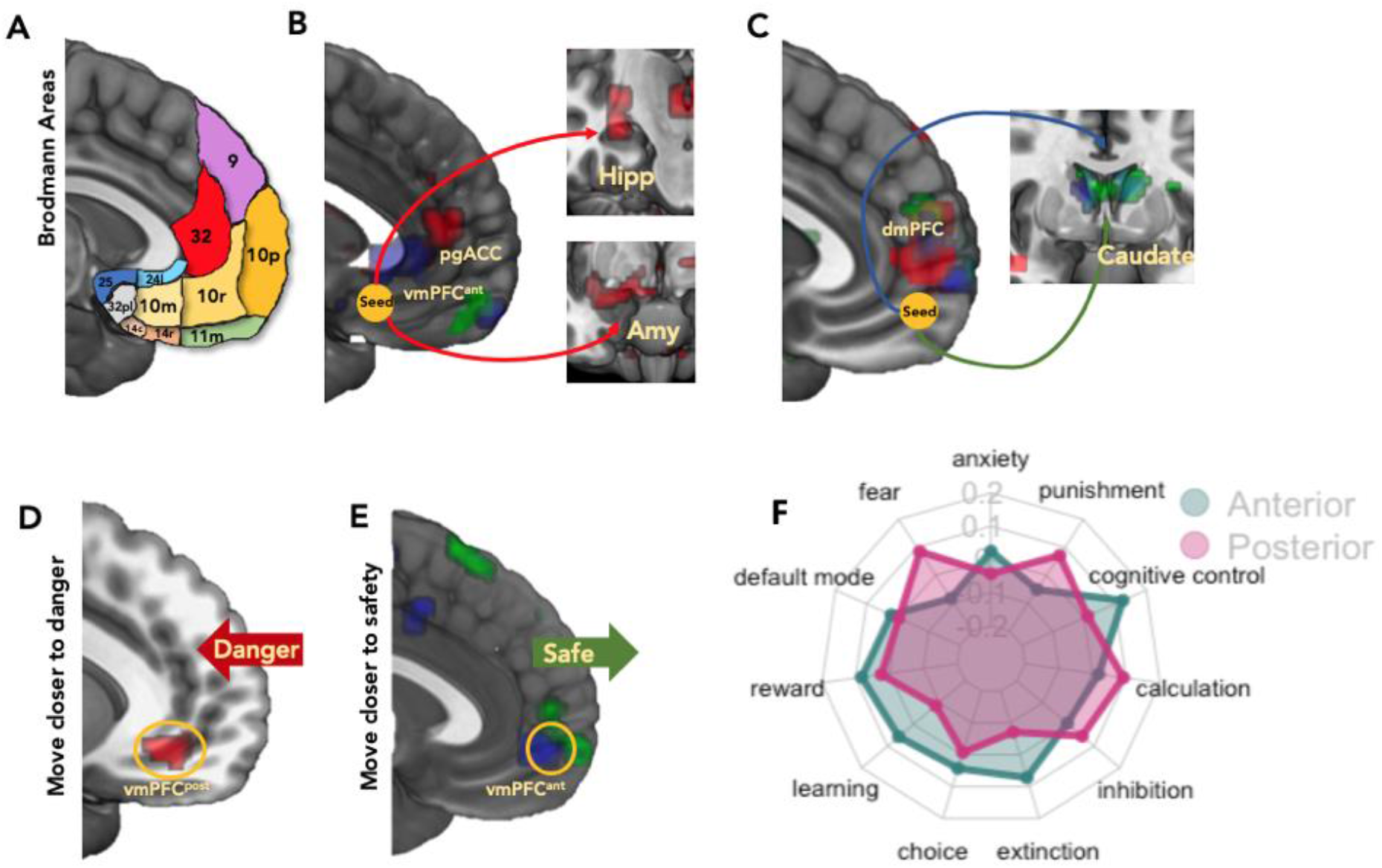
Psychophysiological interactions seeding from regions of interest and meta analytical decoding. (A) Example of Brodmann Areas (BA) that distinguish posterior-anterior axis. For example, the posterior vmPFC reflects BA 25, 24, 32(ACC), 10m and 14, while the anterior encompasses BA 10p, 10 r 11, and 32 (non-ACC). This is made clearer by the dotted line. Connectivity analysis were first performed on the anterior and posterior vmPFC seeds, which are 6 mm spheres centered on the peak voxel of the corresponding clusters in the MVPA searchlight. (B) For the posterior vmPFC seed, in all three attacking conditions, the connectivity maps showed significant activation in the hippocampus (leptokurtic: p < 0.001,T = 4.06; norm_match_: p < 0.001, T = 3.62; norm_half_: p = 0.011, T = 3.18). Interestingly, only in the leptokurtic attacking condition, the amygdala was found significant on the connectivity map (p < 0.001, T = 4.60). (C) On the other hand, with the anterior vmPFC seed, all three attacking conditions showed significant connectivity towards the Caudate (leptokuctic: p < 0.001,T = 3.87; norm_match_ P < 0.001, T = 4.23; norm_half_ P < 0.001, T = 4.59.). We constructed two parametric modulators indicating whether the participants’ final MOS choices is a (D) safety choice or a (E) risky choice (compared to their randomly assigned initial location). The parametric modulation of univariate data thus reveals what regions were associated with risky/ safety choices under different levels of predictability. On the resulting statistical maps, using SVCs from the previously constructed vmPFC_post_ and vmPFC_ant_ ROIs, we found that the “move to danger” and “move to safety” modulations were significant in the vmPFC_post_ and vmPFC_ant_ ROIs respectively (Move to danger: p < 0.001, T = 6.44; Move to safety: p < 0.001, T = 4.39) (F) Meta-analytical decoding with Neurosynth. Red and Green radar bars represent correlation strength between key words and the anterior (x = 0, y = 26, z = −12) and posterior(x = −2, y = 46, z = −10) vmPFC ROIs.

### Differences in vmPFC_post_ and vmPFC_ant_ connectivity

With vmPFC_post_ and vmPFC_ant_ identified as key regions associated with risky and dangerous choices, we were interested in how these regions regulate MOS decisions in concert with subcortical structures. To test this, we performed connectivity analysis using gPPI (see supplementary methods), to reveal regions that showed covarying activity with our vmPFC seed regions. From the MVPA analysis, we took the vmPFC_post_ and vmPFC_ant_ as seed regions for the leptokurtic distribution contrast and normal distribution contrasts, since they were identified as regions representing the process where participants make risk decisions under the corresponding predator conditions. PPI analyses were first performed on the moving to safety/danger contrast, respectively on the vmPFC_post_ and vmPFC_ant_, ROIs (fig 4 b c) For the vmPFC_post_ seed, in all three attacking conditions, the connectivity maps showed significant activation in the hippocampus (leptokurtic: p < 0.001,T = 4.06; norm_match_: p < 0.001, T = 3.62; norm_half_: p = 0.011, T = 3.18). Interestingly, only in the leptokurtic attacking condition did the amygdala show significant coupling with the vmPFC_post_ (p < 0.001, T = 4.60). On the other hand, with the anterior vmPFC seed, all three attacking conditions showed significant connectivity towards the caudate (leptokurtic: p < 0.001,T = 3.87; norm_match_ P < 0.001, T = 4.23; norm_half_ P < 0.001, T = 4.59.). For a full list of activated regions, please refer to supplementary table 2.

### Subjects continually optimize MOS decisions through adaptive learning from trial outcomes

In order to perform effectively on the task, subjects may continually adjust their policy depending on their perceived likelihood of escape which is updated on every trial depending on its outcome. We sought to test this by fitting a simple reinforcement learning model to the behavioral data which assumes subjects estimate the likelihood of receiving a given reward (which depends on both the available reward level and the likelihood of survival) on each trial.

This took the form of a standard Rescorla-Wagner learning model which was used to characterize participants’ margin of safety choice behaviors. The learning rate ‘α’ reflects to what extent participants’ choice of MOS is based on the most recent outcomes. A high learning rate indicates that choice behavior is updated in a more rapid manner based on the difference between the expected choice outcome and the actual choice outcome. In contrast, at low learning rates, surprising outcomes lead to little change in their choice on the next trial. In the current study, we estimated participants’ learning rates in the uncertain vs more certain attack position blocks by fitting a reinforcement learning model36 to their choices in each task block (10 trials per session, as described in figure 1).

We first examined whether our model recapitulated observed patterns in the MOS decision data. The model demonstrated behavior that was consistent with the true data (Figure 5 a), indicating that a reinforcement learning model can describe subjects’ behavior in the task. We next assessed whether participants, as a group, adapted their learning rate in response to the change in attack distances between the more predictable normal distributed attack distances and more uncertain attack distances characterized by leptokurtic outliers. Consistent with previous studies of reinforcement learning, participants’ learning rates were higher in the leptokurtic attack than the more predictable normally distributed attacks positions. (Main effect of attack distribution: F(2,63) = 4.43, p = 0.0159. Post hoc comparisons, p<0.001) (figure 5 b), indicating that subjects adapted their learning based on the level of uncertainty in the attack distribution.

### MOS prediction errors are tracked by a distributed network of brain regions

A parametric modulation analysis on univariate data, using the prediction error from the RL model was performed to address what underlying neural processes were involved during the learning process of participants’ MOS decisions. Small volume corrections were performed on the key ROIs: hippocampus: leptokurtic: p = 0.002; norm_match_: p = 0.004; norm_half_: p = 0.191; amygdala: leptokurtic: P = 0.014; norm_match_: p = 0.006; norm_half_: P = 0.094; striatum: leptokurtic: p < 0.001; norm_match_: p < 0.001; norm_half_: p < 0.001. This suggests that while the striatum decodes the representation of prediction error in all three attacking distributions, the hippocampus and amygdala were involved only in the leptokurtic and norm_match_ attacking conditions. (figure 5 d e).

**Fig. 5.**
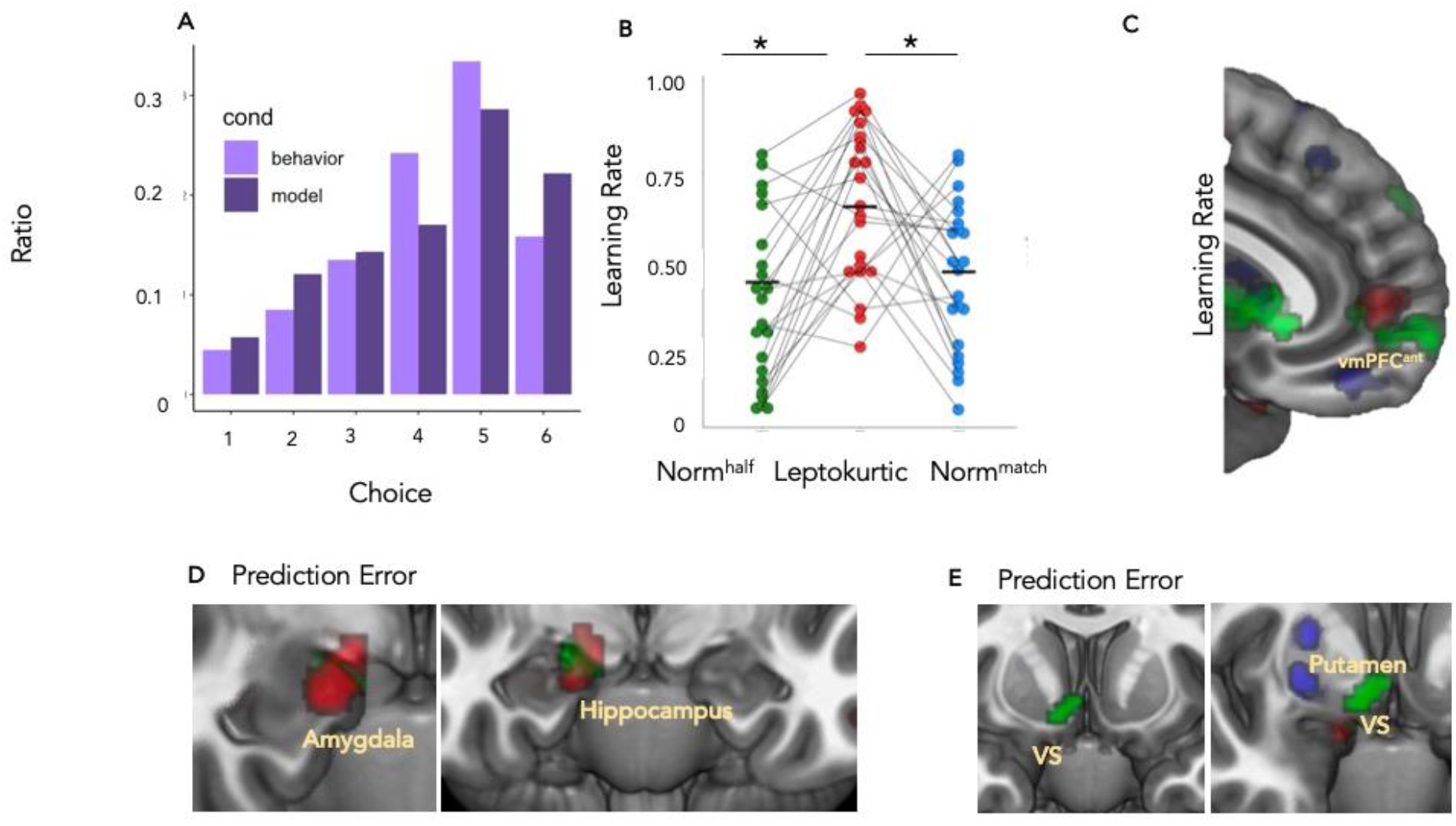
Reinforcement Learning model. (A) Actual MOS choice categories and model fitting MOS choice categories. Choice 1~6 are choice categories from risky to safe. Y axis represents the choice ratio under each category (B) Learning rate from the reinforcement learning model over two days. Data of two sessions within one day were averaged across participants. Learning rate in the leptokurtic condition (which is more predictable) was significantly higher than the other two conditions (posthoc p < 0.001).(C): Maps showing parametric modulation with prediction errors from the model. Small volume corrections (D): (hippocampus): leptokurtic: p = 0.002,; norm1: p = 0.004; norm2: p = 0.191; (amygdala): leptokurtic: P = 0.014; norm1: p = 0.006; norm2: P = 0.094 (E): (striatum): leptokurtic: p < 0.001; norm1: p < 0.001; norm2: p < 0.001. For the remaining activated regions, please refer to supplementary table 3.

## Discussion

We found evidence in support of our hypothesis that in uncertain environments, participants adjust their distance to be closer to safety^37^. We also show that when encountering a more uncertain threat, participants decreased confidence in escape success, while displaying higher learning rates, signifying that under uncertain environments, people adjust decisions more based on recent, immediate information, instead of accumulated information over time. Our MVPA analysis shows that the vmPFC_Post_ is associated with avoidance of more uncertain threats and consequently the decision to stay closer to safety. The vmPFC_Post_ also showed increased functional coupling with the hippocampus and amygdala, supporting the known connectivity with this region as well as its role in control of fear^38,39^. On the other hand, the vmPFC_Ant_ was associated with more certain attack locations and thereby executing safer decisions. These results are congruent with the idea that vmPFC sub-regions play distinct roles in both danger and safety signals that reflect the ability to predict positive or negative outcomes with a threat.

Our results suggest that when the attack location is relatively predictable (i.e. norm_match_ and norm_half_ Guassian distributions), participants make more risky MOS choices. That is, subjects choose to place themselves further away from the safety exit to earn more reward. On the other hand, when the attack location is more unpredictable (i.e. leptokurtic distribution), participants tended to place themselves closer to safety and thus displayed more protective actions. Critically, despite significant differences in variance, there were no differences in MOS decisions between the two Guassian distributions. This suggests that participants’ decision patterns facing uncertain threats was not swayed by a simple change in distribution variance, but by a total structural change in the predictability of the distribution. This was echoed in participants’ subjective rating of their confidence, a reflection of how likely they felt they were to escape (Fig. 2E).

When dissecting the defensive circuitry, it is critical to understand which brain regions are involved in the avoidance of forthcoming danger. Our MVPA searchlight identified three key regions, namely the hippocampus, the vmPFC_Post_ and the vmPFC_Ant_. Interestingly, when looking at the classification accuracies, we found that within the vmPFC_Ant_, classification accuracy was above chance level only for the norm_half_, in line with our prediction that this region would be involved in the most more predictable attack locations. On the other hand, within the vmPFC_Post_, the classification was more accurate than chance level only for the more unpredictable, leptokurtic distribution condition. This suggests a separation of vmPFC subregions in terms of functional roles. While the vmPFC_Ant_ is correlated with more predictable decision environments, the vmPFC_Post_ seems to be associated with more volatile counterparts. Interestingly, the hippocampus classification accuracies revealed no differences between attack locations distributions, suggesting a more general role in avoidance decisions.

The vmPFC_Post_ may function as a hub when the environment is more uncertain and where more information gathering is needed. Further evidence for this comes from our parametric modulation analysis using relative MOS from the starting position, which showed that more dangerous choices are associated with activation in the vmPFC_Post_. This suggests a tentative role for the vmPFC_Post_ to be responsible for computations concerning a more unpredictable environment, or a more risky choice. In our connectivity analysis seeding from the vmPFC_Post_, we observed activations in amygdala and hippocampus only in the uncertain attacking locations. Previous research has shown a role for the amygdala-mPFC as a pathway of modulating threat avoidance behavior^1–4^, and hippocampus as a center for representing predictive relationships between environmental states^17,18^. This is in line with the idea that for decision making under threat with less predictability, more predictive computations are required.

The vmPFC_Ant_ modulates behavior when the environment is relatively easy to predict during the spatial MOS decisions. Interestingly, using relative MOS from the starting position as a modulator in the parametric modulation analysis, the vmPFC_Ant_ was also activated when the choice is categorized as “safe”. In previous studies, this region has been implicated in both safety learning through extinction and safety learning through active avoidance^19,20^. For example, studies using the lever press avoidance task in rodents have shown activation of the prelimbic regions of MPFC (the rodent homologue of human anterior vmPFC) during the expression of active avoidance^43,44^. These regions partially overlap with the identified clusters of vmPFC_Ant_ in our task. Further, when looking at functional connectivity seeding from the vmPFC_Ant_, the caudate was significant only in the two more predictable predator conditions. Although there may be other explanations (action selection^45^). This resonates with previous studies where vmPFC not only functions as a center for signaling safety, but also in reward related processes, because safety processing may be “intrinsically rewarding or reinforcing”^19^. This is also supported by a parametric modulation analysis showing that shifts towards safety activate the vmPFC_Ant_. Also involved in this process is the striatum, which has been shown to be responsible for fear memory extinction^46,47^. For example, previous research on rodents has shown that in rats, the dopamine level in the striatum was unchanged after exposure to novel environmental stimulus, but follows more closely to the expression of conditioned response^28^. Interestingly, this orchestras with our finding where the striatum is only responsive to the high predictability threats together with the vmPFC_Ant_.

We further correlated the neural data with behavioral parameters from the exploratory reinforcement learning model. Parametric modulation using prediction error from the RL model also activated the amygdala in the more uncertain, leptokurtic attacking condition, providing additional evidence for the modulation mechanism where amygdala is involved in the more volatile threat conditions when large discrepancies between expected and observed outcomes happen. Within all predator conditions, the ventral striatum and putamen were also significantly activated in correlation with the PE signal. This is consistent with previous studies where learning under uncertain environments occurs through reward based pathways^48,49^. On the other hand, parametric modulation using learning rates established vmPFC_Ant_ as a hub for MOS decision making when facing predictable attack distances.

The hippocampus also emerged as a central region involved in MOS decisions. First, decoding of choice was higher than chance level in the hippocampus, regardless of how uncertain the attacking locations were. However, the hippocampus only showed functional connectivity with the vmPFC_Post_ in the uncertain, leptokurtic attacking condition. The first finding resonates with the idea that the hippocampus has long been thought of as a predictive map and center for planning when considering future actions based on immediate feedback from the environment^17,18,22^. It was thus universally involved regardless of the uncertainty level of the attacking environment. However, our results indicate that activity in the hippocampus becomes more coordinated with the vmPFC_Post_ in situations which require more intensive planning, as evidenced by the distinct functional connectivity to the hippocampus when the subjects are encountering a more unpredictable, leptokurtic, attacking threat. Indeed, our finding corresponds to previous studies using rodents where the hippocampus has been shown to specifically contribute to model based planning, that may include also memory based decision making^23^.

The current study offers the first insight into how spatial MOS decisions are determined in threat environments with different levels of predictability. It also establishes the posterior and anterior vmPFC subregions as centers modulating the push and pull between risky and safe choices, where the hippocampus is involved in both processes in a more universal manner. More work is needed to further validate the functional separation of vmPFC subregions in terms of their roles during decision making under threat. These new insights, however, suggest a dissociable role of the vmPFC in anxiety, where the vmPFC_Post_ is involved in heightened threat signals, while the vmPFC_Ant_ may be involved in down regulation of threat via safety signals.

## Supplementary Material of

**Supplementary Figure 1:**
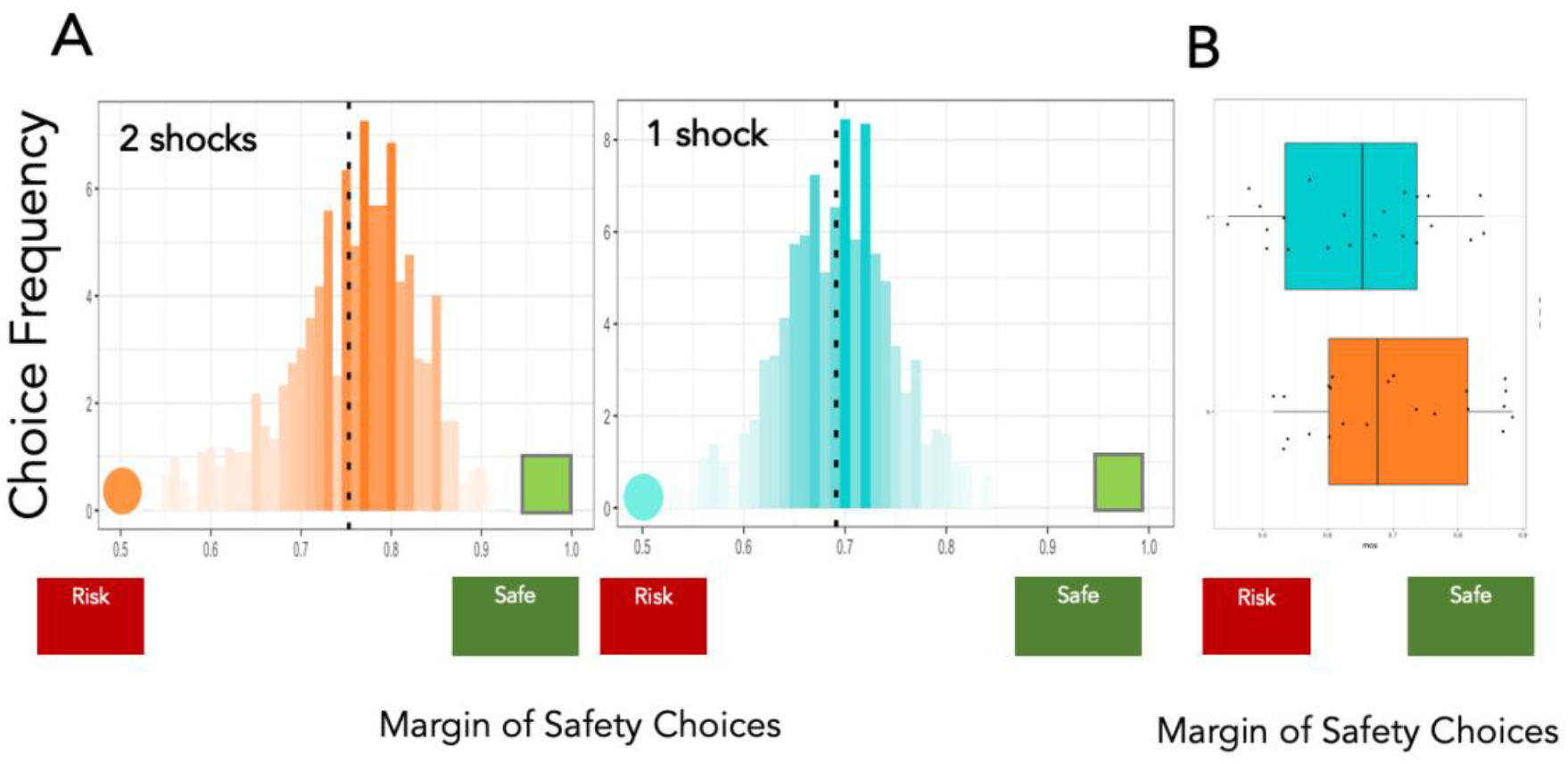
MOS choices under different levels of electrical shocks. (A): X axis represents MOS decision choices, while Y axis represents choice frequency aggregated through all participants and trials. Orange and blue bars deceits conditions from 2-shock and 1-shock respectively. While in the 2-shock condition, participants’ MOS choices were significantly larger than the 1-shock condition (p < 0.001). (B): Same frequency distribution displayed in dot plot.

**Supplementary Figure 2:**
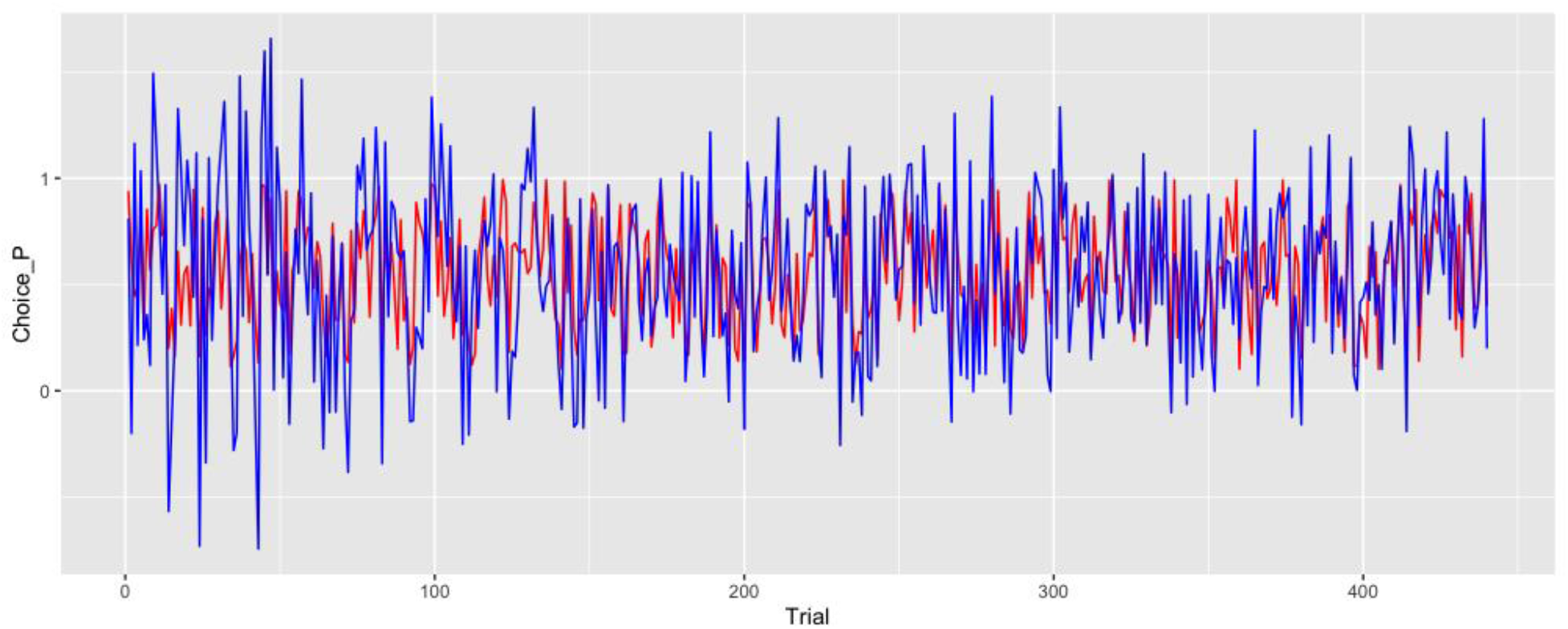
Behavior fitting for the reinforcement learning model. Graph showing the average trajectory of chosen MOS (red line) and model-predicted MOS (blue line) across all participants. X axis represents trial number, while Y axis represents the probability of choosing MOS category 1. It can be observed that fitting improves as trial number increases.

**Table S1.**
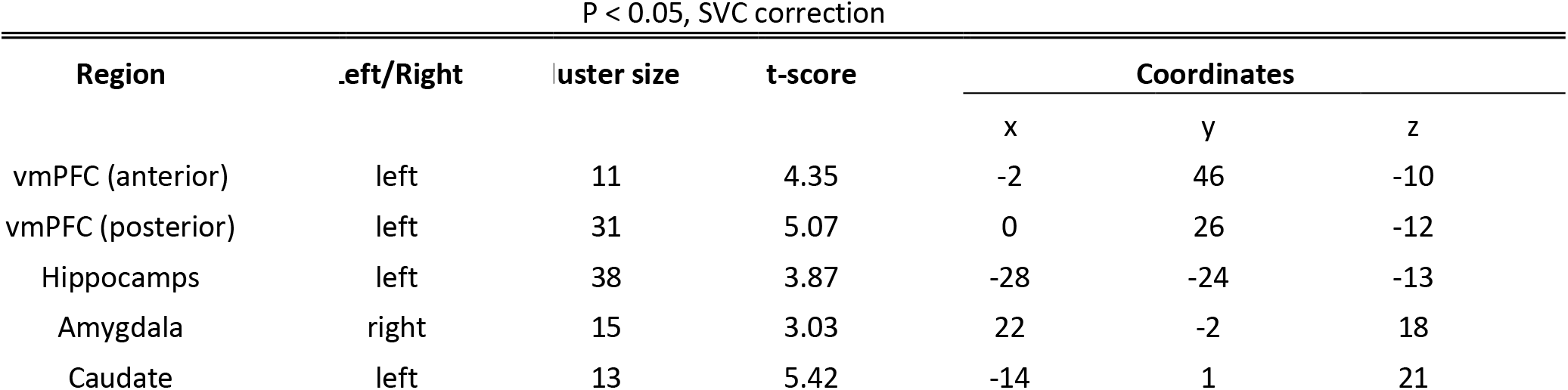
Activation Table for SVC corrections. **Supplementary table 1:** Small volume correction statistics for the ROIs listed in the main paper. Thresholded at P < 0.05 FEW. For the contrasts SVCs, coordinates indicates peak coordinate from the resulting cluster.

| Region | .eft/Right | luster size | t-score | Coordinates |  |  |
| --- | --- | --- | --- | --- | --- | --- |
|  |  |  |  | x | y | z |
| vmPFC (anterior) | left | 11 | 4.35 | -2 | 46 | -10 |
| vmPFC (posterior) | left | 31 | 5.07 | 0 | 26 | -12 |
| Hippocamps | left | 38 | 3.87 | -28 | -24 | -13 |
| Amygdala | right | 15 | 3.03 | 22 | -2 | 18 |
| Caudate | left | 13 | 5.42 | -14 | 1 | 21 |

**Table S2.** Activation Table for PPI analysis. **Supplementary table 2:** psychophysiological Interactions for each predator type, seeding from anterior and posterior vmPFC. Thresholded at P < 0.05 FDR. For the contrasts SVCs, coordinates indicates peak coordinate from the resulting cluster.

|  |  |  |  |  |  |  |
| --- | --- | --- | --- | --- | --- | --- |
| MPFC(anterior) | *ep tokurtic* |  |  |  |  |  |
| **Region** | **.eft/Right** | **luster size** | **t-score** | **x** | **y** | **z** |
| lle Temporal Gyrus | right | 15 | 4.14 | 45 | 0 | -24 |
| usiform Gyrus | right | 14 | 4.98 | 36 | -42 | -18 |
| Lingual Gyrus | left | 43 | 4.21 | -18 | 87 | -9 |
| dial Frontal gyrus | right | 125 | 4.4 | 6 | 57 | 18 |
| MPFC(posterior) | *ep tokurtic* |  |  |  |  |  |
| **Region** | **.eft/Right** | **luster size** | **t-score** | **x** | **y** | **z** |
| Amygdala | right | 32 | 4.95 | 18 | 15 | -3 |
| rior Frontal gyrus | left | 89 | 4.19 | -30 | 57 | -6 |
| usiform Gyrus | left | 26 | 3.38 | -51 | -6 | -27 |
| ior Occipital Gyrus | right | 17 | 4.02 | 48 | -84 | -12 |
| Hippocampus | right | 51 | 4.55 | 21 | -30 | 3 |
| MPFC(anterior) | *mal-matched* |  |  |  |  |  |
| **Region** | **.eft/Right** | **luster size** | **t-score** | **x** | **y** | **z** |
| ior Temporal Gyrus | left | 12 | 4.28 | -33 | 15 | -27 |
| dial Frontal Gyrus | right | 101 | 4.57 | 45 | 42 | -3 |
| dial Frontal Gyrus | right | 224 | 4.45 | 6 | 72 | 0 |
| Thalamus | left | 44 | 4.34 | -18 | 15 | 15 |
| MPFC(posterior) | *mal-matched* |  |  |  |  |  |
| **Region** | **.eft/Right** | **luster size** | **t-score** | **x** | **y** | **z** |
| dial Frontal Gyrus | left | 66 | 3.26 | -3 | 60 | -15 |
| Putamen | right | 35 | 3.63 | 24 | 15 | -9 |
| Precuneus | right | 50 | 4.27 | 3 | -51 | 18 |
| MPFC(anterior) | *ormal-half* |  |  |  |  |  |
| **Region** | **.eft/Right** | **luster size** | **t-score** | **x** | **y** | **z** |
| ior Temporal Gyrus | right | 26 | 4.18 | 33 | 9 | -18 |
| rior Frontal Gyrus | right | 57 | 4.63 | 45 | 45 | -3 |
| dial Frontal Gyrus | right | 69 | 3.98 | 6 | 72 | 0 |
| MPFC(posterior) | *ormal-half* |  |  |  |  |  |
| **Region** | **.eft/Right** | **luster size** | **t-score** | **x** | **y** | **z** |
| Puamen | right | 19 | 4.71 | 21 | 9 | -6 |
| terior Cingulate | left | 24 | 4.99 | 0 | 24 | -3 |
| recentral Gyrus | right | 16 | 4.16 | 39 | -6 | 30 |
| Hippocampus | left | 21 | 3.96 | -20 | -30 | 0 |

**Table S3.** Activation Table for Parametric Modulation with RL Prediction Error. **Supplementary table 3:** Activated regions associated with prediction errors in the RL model. Thresholded at P < 0.05 FDR. For the contrasts SVCs, coordinates indicates peak coordinate from the resulting cluster.

| Region | .eft/Right | luster size | t-score | Coordinates |  |  |
| --- | --- | --- | --- | --- | --- | --- |
|  |  |  |  | x | y | z |
| Fusiform Gyrus | left | 22 | 4.14 | -57 | -21 | -27 |
| rahippocampal Gyrus | right | 45 | 4.431 | 12 | -6 | -15 |
| middle Temporal Gyrus | left | 25 | 4.18 | -63 | -15 | -12 |
| inferior Frontal Gyrus | right | 64 | 4.68 | 63 | 9 | 12 |
| Precentral Gyrus | left | 34 | 3.87 | -60 | 0 | 18 |

**Table S4.**
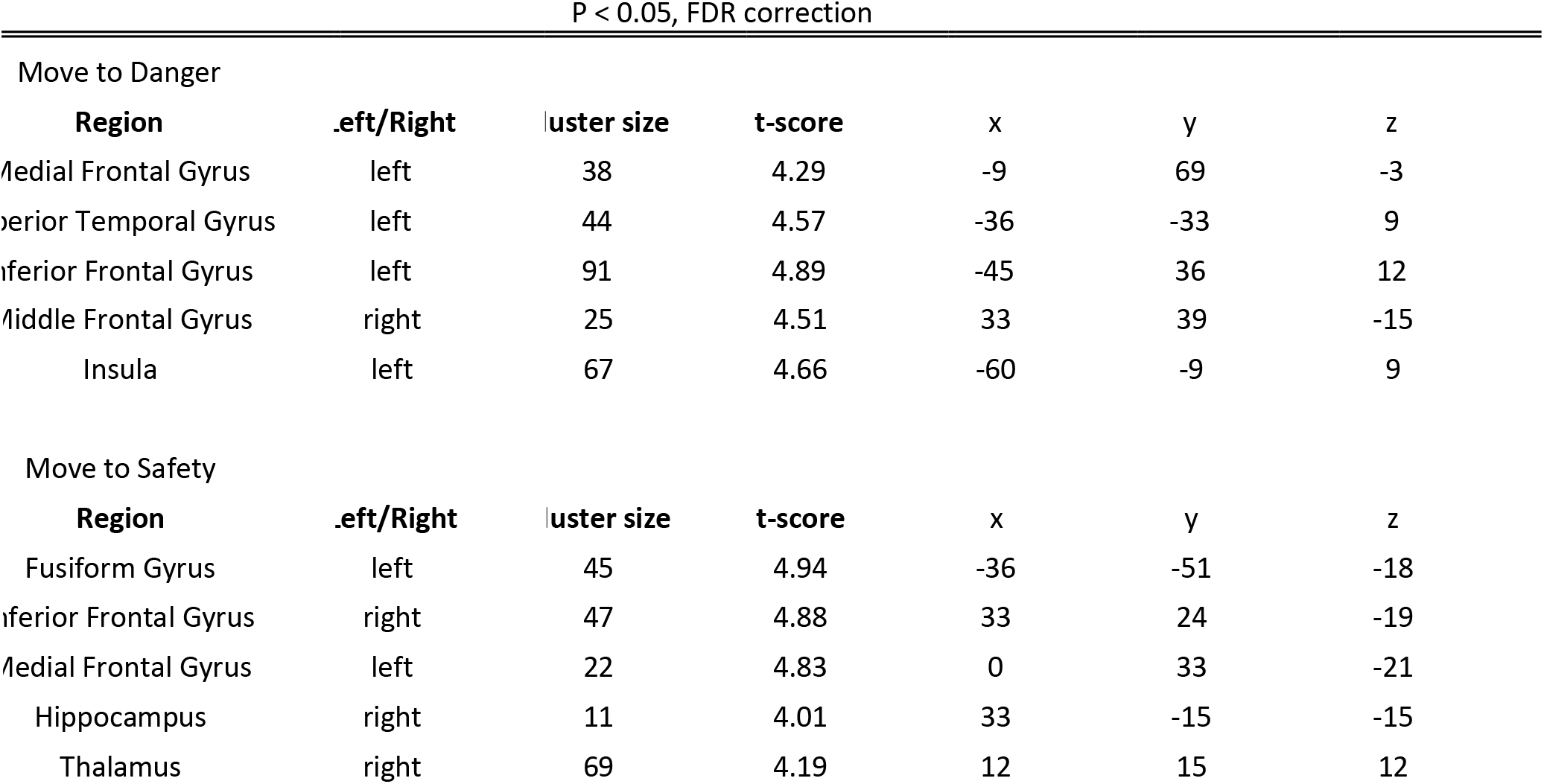
Activation Table for Parametric modulation with escape choices. **Supplementary table 4:** Activated regions associated with parametric modulation of MOS choices (moving to danger or safety). Thresholded at P < 0.05 FDR. For the contrasts SVCs, coordinates indicates peak coordinate from the resulting cluster.

## Supplementary Methods

### Experimental Methods

We tested 24 subjects were recruited according to the guidelines of the California Institute of Technology Institutional Review Board after providing informed consent. Data from two subjects were lost due to incomplete scanning sessions. Our final sample consisted of 22 subjects (10 females, age = 24.3 +−8.1 years).

### Stimuli, apparatus and procedure

A complete pipeline of experimental procedures can be found in figure 1. Participants completed a computer-based task while in an fMRI scanner. The task was set under the scenario where subjects place themselves at a desired location towards a safety exit while facing a potentially dangerous predator. The closer they place themselves to the safety exit, the more likely they will be able to escape from the predator after the trial starts, but the low the resulting reward will be. The goal of the task was to earn as much money as possible while avoiding being caught by the virtual predator. Prior to the beginning of the trial, the participants were presented with a 2 second cue indicating one of the three different predator types that would be presented in the upcoming trial. These predators differ in the location they speed up. These locations correspond to three distributions - a leptokurtic distribution, a normal distribution with matching variance, and a normal distribution with only half of the variance. (In the rest of the paper, we’ll refer to the normal distribution with matching variance as norm_match_, and the normal distribution with half variance as norm_half_) The participants were then shown a two-dimensional runway (90 units distance, where a unit is the smallest increment on the runway), with a triangle icon representing the position of the participant toward the end of the runway (at 80 or 0 units distance, depending on which direction the trial goes. A random starting location is then assigned based on which direction they start), and a circle icon representing the position of a predator at the left side of the runway (at 1 unit distance). This predator had two distinct modes of movement. In ‘‘approach’’ mode, the predator would proceed rightward along the runway at 4 units per second. At a randomly chosen distance (i.e. the attack distance) according to the leptokurtic, norm_match_ and norm_match_ distribution, The predator would switch to “chase’’ mode, at which point it would advance at 10 units per second. The position where it swtiches to the “chase” mode is drawn either from the leptokurtic, norm_match_ or norm_half_ distributions depending on the actual attacking condition.

Before the above mentioned chasing sequence starts, the participants were told to make a decision of where they want to start by pressing left or right arrows, to move from their randomly assigned initial location to a location they desire. The direction of the chase was counter balanced by adjusting the relative location of the predator, participant and the safety zone so that half of the chase was from the left to the right, and the other half were opposite. After participants responded with their preferred margin of safety choice (MOS choice), they skip the actual animation of the chase (which was shown in full during the practice session), and was shown the final result of the trial: whether they got caught or not, and how much reward they earned.

The experiment starts with the subjects being shown that if captured, they will receive 1 or 2 shocks, and high or low reward if they escape (Fig. 1B). They will then be presented with one of three different colored spheres, each representing different attack distributions of the virtual predators. They will then be asked to rate how confident they are of escape. Next, the subject will be asked to make safety decisions by either staying or switching to a riskier position that is further away from the safety exit or stay or move closer to the safety exit. To motivate risky decisions, the subject will acquire more money if they are more risky (i.e. move further from safety), which follows a simple linear relationship as a function of MOS choice (10 cents minimum, 2o cents maximum). They will then be asked to move the cursor to the decided safety position. After a jittered ITI, the subject will observe the outcome. If caught, they will receive a shock(s) and lose their money on this trial. This will repeat for another nine trials, before the subject is introduced to a new set of reward and shock contingences as well as a new virtual predator. The virtual predator attack distribution is either (i) normal distribution with half variance, (ii) leptokurtic (positive kurtosis with fatter tails) or (iii) normal distribution with matched variance with the leptokurtic distribution. Leptokurtic distributions are rare in the natural environment, where distributions are often normally distributed and easier to learn.

A total number of 460 trials (400 experimental trials and 60 control trials) were administrated throughout 4 sessions (2 sessions per day with two days). The computer task was programmed in Pygames with Python.

#### fMRI data acquisition

We will collect the fMRI images using a 3T Prisma scanner in the Caltech Brain Imaging Center (Pasadena, CA) with a 32-channel head receive array. BOLD contrast images will be acquired using a single-shot, multiband T2*-weighted echo planar imaging sequence with the following parameters: TR/TE = 1000/30 ms, Flip Angle = 60°, 72 slices, slice angulation = 20° to transverse, multiband acceleration = 6, no in-plane acceleration, 3/4 partial Fourier acquisition, slice thickness/gap = 2.0/0.0 mm, FOV = 192 mm × 192 mm, matrix = 96 × 96). Anatomical reference imaging will employ 0.9 mm isotropic resolution 3D T1w MEMP-RAGE (TR/TI/TE = 2550/1150/1.3, 3.1, 4.0, 6.9 ms, FOV = 230 m x 230 mm) and 3D T2w SPACE sequences (TR/TE = 3200/564 ms, FOV = 230 mm x 230 mm). Participants viewed the screen via a mirror mounted on the head coil, and a pillow and foam cushions were placed inside the coil to minimize head movement. Electric stimulation was delivered using a BIOPAC STM100C.

### Data Analysis

All statistical analyses for the behavioral data were carried out in R, using the packages ‘ezANOVA’, ‘coxme’, and ‘lme4’. Where appropriate, Greenhouse–Geisser corrections were performed to account for violations of sphericity, and the correction factor values ($\epsilon$) and original degrees of freedom are reported. Partial eta-squared effect sizes are reported only for significant analyses. Where appropriate, we corrected for multiple comparisons using Holm-Bonferroni.

Analysis of fMRI data was carried out using scripted batches in SPM8 software (Welcome Trust Centre for Neuroimaging, London, UK) implemented in Matlab 7 (The MathWorks Inc., Natick MA). Structural images were subjected to the unified segmentation algorithm implemented in SPM8, yielding discrete cosine transform spatial warping coefficients used to normalize each individual’s data into MNI space. Functional data were first corrected for slice timing difference, and subsequently realigned to account for head movements. Normalized data were finally smoothed with a 6-mm FWHM Gaussian kernel.

A multivariate pattern analysis was performed using PyMVPA (Hanke et al., 2009). We extracted the beta values associated with experimental conditions of all the voxels in each ROI, removing the mean intensity for each multi-voxel activity pattern. For each participant, the brain response pattern analyses of classification training and testing with linear support vector machines (SVMs) were conducted using a leave-one-run-out cross-validation procedure. Furthermore, to evaluate whether stimulus contrast modulates brain response patterns, cross-validations that use low-contrast condition data for training and high-contrast condition data for testing, and vice versa, were also conducted. ANOVAs were then conducted to compare classification accuracies.

To localize the functional ROIs, a whole-brain searchlight was first performed to identify brain regions sensitive to MOS decision information, where a classifier predicting each trial’s association with one of the six MOS decision category was constructed. For each voxel in native space, we built a spherical region of interest (ROI, radius 6 mm) centering on the voxel, extracted t values in this ROI to each of the 50 MOS decisions and calculated one minus Spearman rank correlations of all decision pairs within this ROI to construct a neural RDM. The relationship between the neural RDM and the theoretical RDM was then assessed using partial Spearman correlation, which produced a correlation coefficient for this voxel. Moving the searchlight center throughout the cortex, we obtained a whole-brain r-map in the native space. Note that the searchlight analysis was restricted to the voxels with a probability higher than 1/3 in the native gray matter image generated from the segmentation step. For a group-level random-effects analysis, the r maps in the native space were Fisher-z-transformed, normalized to the MNI space using the forward deformation field and spatially smoothed using a 6 mm full-width at half maximum Gaussian kernel. Clusters surviving the cluster-level FWE correction at P < 0.05 were reported. For each subject, we then identified the voxels whose neural RDMs showed a significantly positive correlation with the RDM in the above-mentioned searchlight analysis (P < 0.05, FDR corrected). These voxels together with their adjacent voxels within a 6-mm-radius sphere were considered as individual subjects’ functional ROI. (figure 3 b c d e).

### Classification accuracy

To explore the regions involved in the decision making process under threat within the Margin of Safety framework, we examined MVPA classification accuracies using both whole brain searchlight analysis and ROI analysis. We extracted voxel-wise fMRI responses to margin of safety trial (decision phase) as classification samples. For each participant and each run, we designed a general linear model (GLM). The GLM contained 3 regressors indicating the decision phases (duration = 4 s) of the 3 distribution types, as well as 4 regressors indicating the indication phase (duration = reaction time), motor phase (duration = 4 s), and feedback phase (duration = 3 s). All the regressors were convolved with a canonical hemodynamic response function. In addition, six motion-correction parameters and the linear trend were included as regressors of no interest to account for motion-related artifacts. For each voxel, the parameter estimates of the 3 regressors corresponded to the fMRI responses to each of the 3 distributions in each run. The fMRI responses to each distribution item were then entered into the classification analysis as classification samples.

Naturally, there are two main questions we prioritized. First, what brain regions are involved in determining which distribution type the participant is facing and second, what brain regions are involved in determining the MOS decision the participant is making. Thus, we used two sets of classification labels corresponding to the two questions: i) norm_match_, norm_half_, and leptokurtic distribution 2) the 50 possible discrete MOS choice options.

We employed a linear support vector machine with a cost parameter C = 1 as a classifier. Classification accuracy was estimated using a leave-one-run-out cross-validation: for each of the four runs, a classifier was trained on the other three runs and tested on the remaining focal run; and the procedure was repeated for the four runs (accuracy scores were averaged).

To validate whether the classification performance was significantly above chance, we further conducted Monte Carlo permutation-based statistical tests. This method entailed running a classification analysis 1000 times with randomly permuted experimental condition labels, allowing us to construct null distributions that were used to examine whether a classification accuracy was significantly above chance at an α of p < 0.05.

### Univariate analyses

We also ran a univariate analysis pipeline to decompose the neural circuits employed when facing uncertain and stable threats. Preprocessed imaged were subjected to a two-level general linear model using SPM8. The first level contained the following regressors of interest, each convolved with the canonical two gamma hemodynamic response function: a 2-second box-car function for the onset of the trial (where the color of the incoming attack is shown); a 4 second (duration jittered) box-car function for the decision period; a 4-second boxcar (function for the time window where participants actually select their starting positions. Mean-centered trait anxiety ratings, and parameters in the reinforcement learning model were included as orthogonal regressors. In addition, regressors of no interest consisted of motion parameters determined during preprocessing, their first temporal derivative and discrete cosine transform-based temporal low frequency drift regressors with a cutoff of 192-seconds.

Beta maps were used to create linear contrast maps, which were then subjected to second-level, random-effects one-sample t tests. In Addition, A flexible factorial model was used to examine the main effects of attack type, reward level and shock level. Interaction effects between attack type, reward level and shock level were also examined using the factorial model. The resulting statistical maps were thresholded at P < 0.05 corrected for multiple comparisons (false discovery rate [FDR] corrected).

### Connectivity Analysis

Based on the key regions obtained during MVPA searchlight analysis, we further performed connectivity analysis using gPPI (gPPI; http://www.nitrc.org/projects/gppi), which is configured to automatically accommodate more than two task conditions in the same PPI model by spanning the entire experimental space, compares to the standard implementation in SPM8. The connectivity model is based on the underlying concept using the following models:

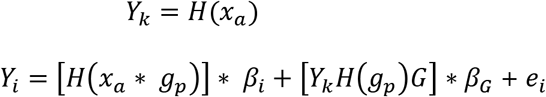

where H is the HRF in Toeplitz matrix form; Yk is the BOLD signal observed in the seed region; xa is the estimated neural activity from the BOLD signal in the seed region (Gitelman et al., 2003); Yi is the BOLD signal observed at each voxel in the brain; βi is a matrix of the beta estimates of the psychophysiological interaction terms; βG is a matrix of the beta estimates of the seed region BOLD signal (Yk), covariates of no interest (G), and task regressors that are the convolution of psychological vectors H(gp); and ei is a vector of the residuals of the model. In the gPPI approach, gp is a vector formed by multiplying the condition ON times (onset times plus stimulus duration — when the stimulus or psychological state is presented to the participant or when the participant experiences a defined psychological/experimental state) by a weighting vector.

